# The potential of aquatic haematophagous, liquidosomatophagous and macrophagous leeches as a tool for iDNA characterisation

**DOI:** 10.1101/2021.05.01.442262

**Authors:** Christina Lynggaard, Alejandro Oceguera-Figueroa, Sebastian Kvist, M Thomas P Gilbert, Kristine Bohmann

## Abstract

Leeches play important roles in food webs due to their abundance, diversity and feeding habits. Studies using invertebrate-derived DNA (iDNA) extracted from leech gut contents to target vertebrate DNA have focused on the Indo-Pacific region and mainly leveraged the leech family Haemadipsidae, composed of haematophagous terrestrial leeches, while the aquatic haematophagous, liquidosomatophagous and macrophagous counterparts have largely been disregarded. While there is general knowledge regarding the taxonomic groups that leeches prefer to feed on, detailed taxonomic resolution is still missing and therefore, their potential use for monitoring animals is not known. In this study, 116 non-haemadipsid leeches belonging to 12 species and spanning the three feeding habits were collected in Mexico and Canada. We used DNA metabarcoding to investigate their diet and assess their potential use for vertebrate monitoring. We detected vertebrate taxa from five orders including fish, turtles and birds in the diet of the aquatic haematophagous leeches; eight invertebrate orders of annelids, arthropods and molluscs in the liquidosomatophagous leeches; and ten orders of invertebrates belonging to Arthropoda and Annelida, as well as one vertebrate and one parasitic nematode, in the macrophagous leeches. These results show the potential use of iDNA from the gut content of aquatic haematophagous leeches for retrieving vertebrate taxa, and from macrophagous and liquidosomatophagous counterparts for invertebrates. Our study provides information about the dietary range of the freshwater leeches and the non-haemadipsid terrestrial leech and proof-of-concept for the use of non-haemadipsid leeches for animal monitoring, expanding our knowledge of the use of iDNA from leech gut contents to North America.

## 1. Introduction

Leeches (Annelida: Hirudinea) are found on all continents including the oceanic waters of Antarctica (Sket & Trontelj, 2008). There are around 680 extant leech species (Sket & Trontelj, 2008) and these have varying and important roles in food webs; they can be ectoparasitic (or, on occasion, endoparasitic [Mann & Tyler, 1963]), predatory, intermediate and final hosts for parasites (Sawyer, 1986), vectors of hemogregarine and trypanosome blood parasites (Barta & Desser, 1989; Siddall & Desser, 1991; Siddall & Desser, 2001) and serve as the primary diet for several fish species across the globe (e.g. Sawyer, 1986; Young & Spelling, 1986). Leeches are also found in terrestrial and marine habitats, yet most of the known leeches inhabit freshwater ecosystems (Sawyer, 1986; Sket & Trontelj, 2008).

Besides haematophagy (parasitism), from now on referred as bloodfeeding, the feeding habits of leeches range from macrophagy (i.e., predation of invertebrates) to liquidosomatophagy (i.e., feeding on internal liquids and soft tissues of invertebrates, mainly molluscs and oligochaetes) (Sawyer, 1986). While there is general knowledge regarding the overall taxonomic groups that leeches prefer to feed on, more detailed taxonomic resolution of the diet of most leeches is still missing. For example, in macrophagous leeches, members of the freshwater families Erpobdellidae and Haemopidae are known to be predators that ingest oligochaetes, other invertebrates and even other hirudineans including members of their own species (Darabi & Malek, 2011; Kutschera & Wirtz, 2001; Simon & Barnes, 1996).

However, information is lacking about preference to a specific taxonomic group, and the prevalence of cannibalism. The paucity of dietary information extends to aquatic bloodfeeding leeches, such as those of the family Glossiphoniidae and members of Macrobdellidae, which are known to feed on blood of vertebrates, but whose specific dietary preferences are largely unknown. Whereas members of the genus *Placobdella* (Glossiphoniidae) are primarily considered parasites of turtles (Sawyer, 1972), some species have been shown to feed on amphibians and birds, and some will readily feed on humans (personal observation). Species of the genus *Haementeria* are known to be parasites of mammals but other vertebrate hosts have been recorded as prey of these leeches, indicating a flexible diet. One species in the family Macrobdellidae, *Macrobdella decora* (Say, 1824) has been found to feed on vertebrate blood, but also preying on amphibian eggs (Trauth & Neal, 2004; Turbeville & Briggler, 2003). More information is needed to ascertain if this behaviour is opportunistic, or a preference. Similarly, members of the family Piscicolidae seem to show a preference for fish blood, including both Chondrichthyes and Osteichthyes; however, some isolated records of leeches feeding on molluscs and crustaceans have been reported (López-Peraza et al., 2017; Nakano, 2017) indicating a broader dietary range. For terrestrial leeches detailed information is also lacking, or is even completely absent such as for the mexican terrestrial leech *Diestecostoma mexicana*, except for the family Haemadipsidae for which the diet is well studied as part of invertebrate derived DNA, or iDNA studies, which aim to monitor vertebrates through DNA detection in leech bloodmeals (Drinkwater et al., 2020; Fahmyet al., 2019; Nguyen et al., 2021; Schnell et al., 2018; Tessler et al., 2018).

Traditional methods for studying the diet of leeches have relied on direct observations of the leeches feeding on other animals (Darabi & Malek, 2011; Kutschera, 2003), or examining the gut content (Sawyer, 2019; Toman & Dall, 1997). However, diet information can be difficult to obtain when working with free-ranging leeches that are not caught while feeding and that have already partially digested the ingested food, or when dealing with macrophagous and liquidosomatophagous leech taxa as the diets will lack morphological characteristics. This has led to leech diets being studied under experimental settings (Darabi & Malek, 2011; Gaudry et al., 2010). In recent years, the use of molecular methods has improved our knowledge on the role of leeches in trophic networks and has greatly increased our understanding of their diets. The emerging molecular field of iDNA has contributed to the knowledge on leech diet and leech-derived iDNA is now used as a complementary tool to traditional vertebrate monitoring methods (Schnell et al., 2012; Ji et al., 2020). Such leech-derived iDNA studies have focused almost exclusively on terrestrial bloodfeeding leeches, and in line with the geographical distribution of (Borda et al., 2008), they have been used to detect vertebrates in Asia (Vietnam, Laos, Malaysia, Japan), Africa (Madagascar) and Oceania (Australia and Tasmania) (Abrams et al., 2019; Drinkwater et al., 2018; Fahmy et al., 2019; Ji et al., 2020; Morishima et al., 2020; Nguyen et al., 2021; Schnell et al., 2018; Schnell et al., 2012; Tilker et al., 2020. As these haemadipsid leeches have an Indo-Pacific distribution, if iDNA from leeches is to be used for vertebrate monitoring elsewhere in the world, the vertebrate dietary range of non-haemadipsid leeches will need to be assessed, i.e. freshwater bloodfeeding, liquidosomatophaous and macrophagous leeches. Only two previous iDNA studies have investigated iDNA from freshwater bloodfeeding leeches. The first study focused on a single leech species (*Haementeria acuecueyetzin*) that could be inferred to actively feed on the Antillean manatee (*Trichechus manatus*) in Mexico, through sequencing of the leech bloodmeal (Pérez-Flores et al., 2016). The second study took a more all-encompassing approach to understanding the diet of the European medicinal leech (*Hirudo medicinalis*) and also leveraged this diet information to pinpoint the geographic area where the leeches had been collected (Williams et al., 2020). Larger scale assessments of vertebrate taxa in the diet of freshwater bloodfeeding leeches, and leeches with other feeding habits, have not been carried out to assess the potential of freshwater leeches in iDNA vertebrate monitoring. In this study, we aim to i) provide insights into the dietary range of aquatic bloodfeeding, liquidosomatophaous, and macrophagous leeches collected in Mexico and Canada, in addition to elucidating the diet of a terrestrial leech *Diestecostoma mexicana*, and use it to ii) assess the potential use of iDNA from these leeches as vertebrate monitoring tools. To achieve this, we applied metabarcoding to DNA extracted from the gut contents of freshwater and non-haemadipsid terrestrial leeches collected in Mexico and Canada. The analyzed leeches span six families and all three feeding modes; bloodfeeding, macrophagy and liquidosomatophagy.

## 2. Methods

### 2.1 Sample collection

A total of 116 leech specimens (Clitellata: Hirudinea) from the families Glossiphoniidae, Piscicolidae, Erpobdellidae, Macrobdellidae, Haemopidae and Xerobdellidae were collected in 2015 and 2018 in different localities in Mexico and Canada. Leech specimens were selected based on their phylogenetic position in order to include representatives of all major lineages. In Mexico, leeches were collected from eight freshwater localities and one terrestrial forest habitat (Table 1, Supplementary information). In Canada, leeches were collected from two separate freshwater localities in Ontario. The collection methods varied for leeches caught in different habitats and with different feeding habits. Non-bloodfeeding freshwater leeches were hand collected from plants, logs and other submerged debris and the terrestrial leech, *Diestecostoma mexicana*, from under rocks on soil. Bloodfeeding freshwater leeches were collected directly from their host (in the case of *Myzobdella patzcuarensis*), from under rocks immersed in water and by submerging the collector’s legs into the water and removing any attached leeches. Importantly, in every case, bloodfeeding leeches were collected before feeding on the collector’s blood (but see Results for one case of human DNA inside of a single leech).

**Table 1.**
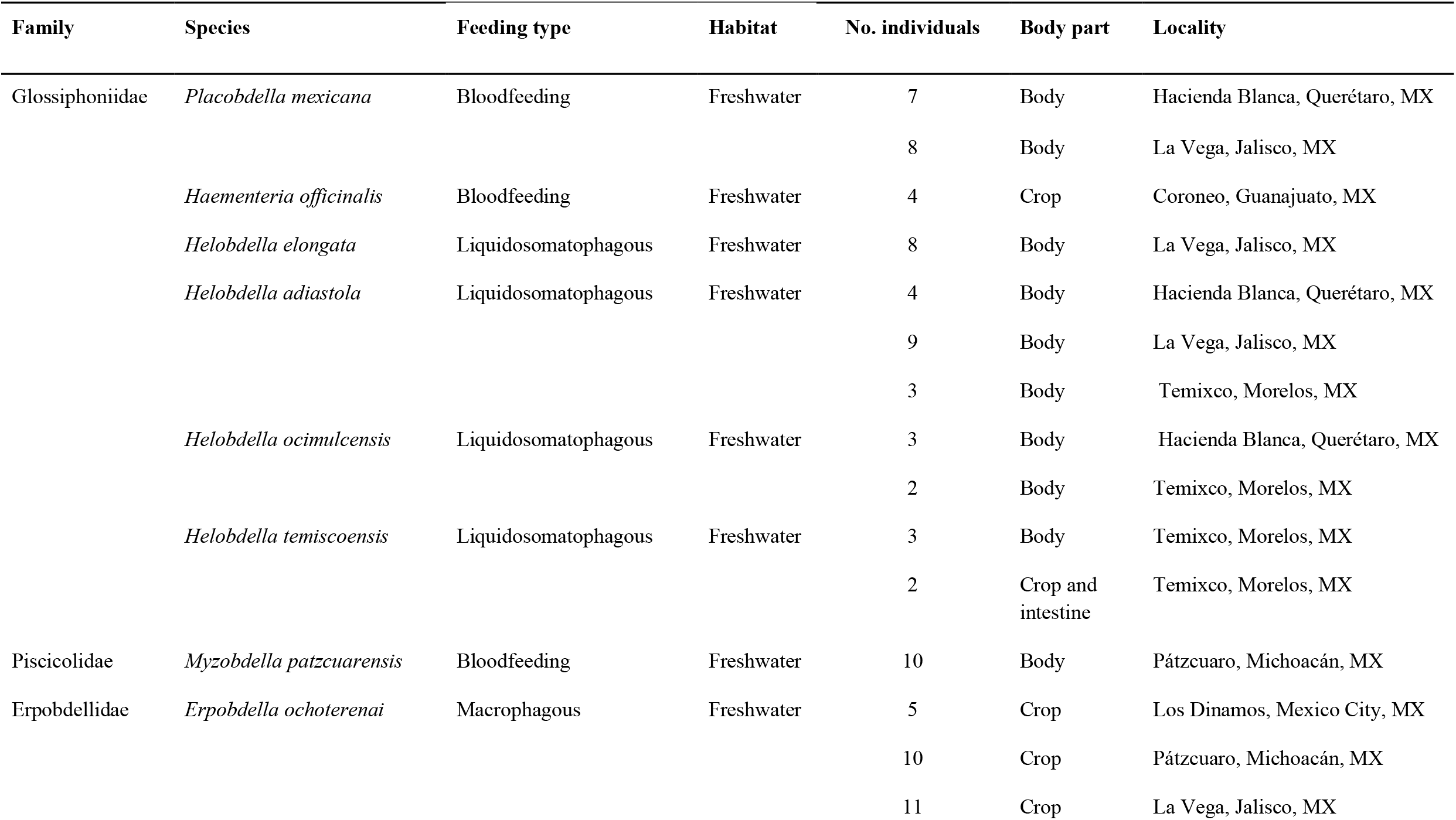
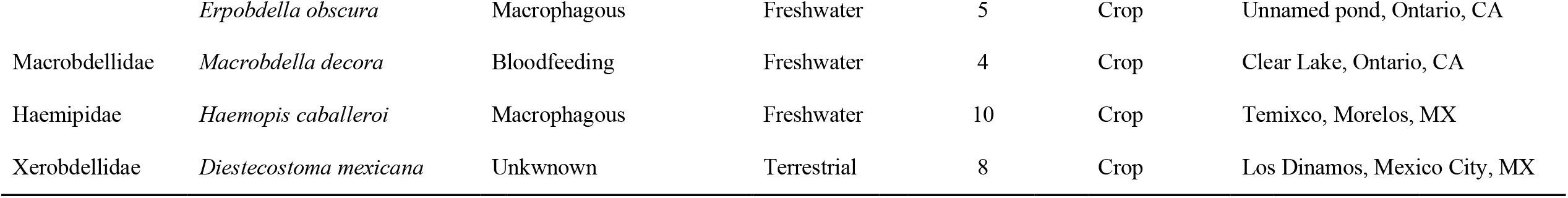
Leeches collected in Mexico (MX) and Canada (CA) grouped by family, and with information regarding their feeding type, the habitat in which they live, number of individuals collected, body part used for DNA extraction and the locality where they were collected.

Prior to DNA extraction, leeches were identified using specialized literature (Klemm, 1985; Sawyer, 1986; Oceguera-Figueroa, 2020). Leeches were sterilised with a 10% bleach solution after both ends of the body were closed with tweezers to avoid bleach entering the leech, and then rinsed with distilled water and stored individually in absolute ethanol in 2 ml Eppendorf or 15 ml Falcon tubes. Specimens were dissected under a stereomicroscope for the removal of the crop and intestine. However, due to their small size, for two individuals of *Helobdella temiscoensis*, the crop and intestine could not be separated so the entire digestive tract was removed as one piece; for the remaining *Helobdella* species and two specimens of *Placobdella mexicana* from Hacienda Blanca, no dissection was possible so the entire body was used for DNA extraction (see Table 1). When dissecting *Erpobdella obscura*, larval nematodes were found encysted in the gastrointestinal tissue, i.e., not in the intestine itself, and were therefore removed and stored in 70% ethanol for further analyses. The entire digestive tracts and complete leeches were kept in absolute ethanol at -20°C until further DNA extraction.

### 2.2. DNA metabarcoding

DNA was extracted using the spin-column protocol for animal tissues from the DNeasy Blood & Tissue Kit (Qiagen, USA) according to manufacturer’s instructions with slight modifications. Firstly, the samples were subjected to three freeze-heat rounds of -80°C for 15 min and 50°C for 30 min prior to addition of Proteinase K. This was done in order to increase the rupture of bacterial membranes (Lever et al., 2015), in the anticipation that the DNA extracts might be used in future microbiome analyses. Secondly, to increase DNA yield, an incubation step of 37°C for 15 mins was added after the addition of 100μl of the elution buffer. A negative extraction control was included for every 20 samples.

The mlCOIintF (forward 5’-GGWACWGGWTGAACWGTWTAYCCYCC -3’) and jgHCO2198 (reverse 5’-TAIACYTCIGGRTGICCRAARAAYCA-3’) metabarcoding primer set was used to PCR amplify ca. 313 basepairs of the mitochondrial cytochrome *c* oxidase subunit I (COI) barcode marker (Geller et al., 2013; Leray et al., 2013), as it is a universal eukaryote primer and expected to amplify both diet and host DNA, thereby enabling molecular verifications of the leech identities. To allow multiplexing, nucleotide tags were added to the 5’ ends of both forward and reverse primers (Binladen et al., 2007). Specifically, tags consisted of a total of 7-8 nucleotides of which 6 nucleotides were nucleotide tags and 1-2 were nucleotides added to increase complexity on the flow cell during sequencing (De Barba et al., 2014).

Prior to the metabarcoding PCR amplifications, a dilution series (1:5 and 1:10) of a subset of the DNA extracts, and positive and negative controls, were screened using SYBR Green qPCR with the aim to determine optimal cycle number for the subsequent PCRs, screen for contamination in the negative controls and in the samples, and calculate the maximum DNA template in which PCR inhibitors would not distort amplification. The 20 μL reaction consisted of 1μL DNA template, 1μL of SYBR Green/ROX solution [one part SYBR Green I nucleic acid gel stain (S7563) (Invitrogen), four parts ROX Reference Dye (12223-012) (Invitrogen) and 2000 parts high-grade DMSO], 0.75 U AmpliTaq Gold, 1x Gold PCR Buffer and 2.5 mM MgCl_2_ (all from Applied Biosystems), 0.6 μM each of 5’ nucleotide tagged forward and reverse primers, 0.2 mM dNTP mix (Invitrogen) and 0.5 mg/mL bovine serum albumin (BSA). The thermocycling profile was 95°C for 10 min, followed by 40 cycles of 95°C for 15 seconds, 51°C for 30 seconds and 72°C for 60 seconds, followed by a dissociation curve. The amplification plots indicated that the use of a 1:5 diluted DNA extract and running 27 cycles was optimal across samples to be used in the following metabarcoding PCR amplification.

Tagged PCRs were subjected to three PCR replicates for each of the 116 DNA extracts, all negative extraction controls and two positive controls [DNA extracted from lion (*Panthera leo)* and giraffe (*Giraffa camelopardalis*)]. Further, negative controls were included for every seven DNA extracts. PCR amplifications were performed with non-matching nucleotide tags (e.g. F1-R2, F1-R3, F1-R4…) to allow for more amplicons to be pooled together and reduce laboratory costs (Schnell, Bohmann, & Gilbert, 2015). The 20 μL reactions were set up as described for the qPCR above but omitting SYBR Green/ROX and replacing the dissociation curve with a final extension time of 72°C for 7 min. Amplified PCR products were visualized on 2% agarose gels with GelRed against a 100 bp ladder. All negative controls appeared negative and all DNA extracts and positive controls showed successful amplification. PCR products of DNA extracts, including negative and positive controls carrying different nucleotide tag combinations, were pooled resulting in three amplicon pools: one pool per replicate.

Amplicon pools were purified with MagBio HiPrep beads (LabLife) using a 1.6x bead to amplicon pool ratio and eluted in 35 μL EB buffer (Qiagen). Purified amplicon pools were built into sequence libraries with the TagSteady protocol to avoid tag-jumping (Carøe & Bohmann, 2020). Libraries were purified with a 0.8x bead to library ratio and eluted in 30μL EB buffer and qPCR quantified using the NEBNext Library Quant Kit for Illumina (New England BioLabs Inc.). Purified libraries were pooled equimolarly according to the qPCR results and sequenced at the GeoGenetics Sequencing Core, University of Copenhagen, Denmark. Libraries were sequenced using 250 bp paired-end reads on an Illumina MiSeq sequencing platform using v2 chemistry, aiming at 25,000 paired reads per PCR replicate.

### 2.3. Data processing

Illumina adapters and low quality reads were removed, and paired reads with a “minalignment” score of 50 and “minlength” score of 100 were merged using AdapterRemoval v2.2.2 (Schubert et al., 2016). Sequences were sorted within each amplicon library based on primer and tag sequences using Begum (https://github.com/shyamsg/Begum), allowing for two mismatches to primer sequences and no mismatches to tag sequences.

Begum was further used to filter sequences across the three PCR replicates for each sample. Filtering parameters were set according to the sequenced negative and positive controls and sequences present in a minimum of two out of three PCR replicates and with a minimum of 25 copies were retained. Sequences were clustered into operational taxonomic units (OTUs) with 97% similarity using SUMACLUST (https://git.metabarcoding.org/obitools/sumaclust/wikis/home/). The LULU algorithm (Frøslev et al., 2017) was used with default settings to detect and remove erroneous OTUs. None of the OTUs found in the positive or negative controls were found in the samples, indicating that there were no tag-jumps (Schnell, Bohmann, & Gilbert, 2015) or cross contamination.

Taxonomic assignment to the OTU sequences, including the leeches and the gut contents, was performed using BLASTn against the NCBI non-redundant (nr) sequence database (www.ncbi.nlm.nih.gov/), and the output was imported into MEGAN Community Edition v6.12.7 (Huson et al., 2016) using a weighted LCA algorithm with 80% coverage, top percent of 10, and a min score of 150. Genus and species information was complemented with data retrieved from BOLD (Barcode of Life Database, http://www.boldsystems.org/). OTUs were assigned to species-level taxa if they had a percentage of identity higher than 99% to a reference sequence and matching to only one species. It should be noted that, during dissection, entire leeches were found in the crop of *Erpobdella* species, including individuals of their own species. Because of this, if detecting more than one OTU assigned to *Erpobdella*, OTUs with the highest number of reads were considered as sequences belonging to the predator, and the remainder were considered gut contents. OTUs that could not be identified to any taxonomic level were discarded from further analyses.

## 3. Results

A total of 10,209,408 raw sequence reads were generated from the amplicon libraries. After filtering steps 3,824,526 reads were retained for downstream analyses, and a total of 69 OTUs were detected in the analysed leech gut contents. The number of reads after each filtering step can be found in Supplementary Table 1. In addition to the identification of the leech taxa, the identified taxa included vertebrate and invertebrate taxa. In all cases, the morphological identifications of the analysed leeches were confirmed by the DNA analysis.

### 3.1 Animal taxa detected in leech gut content

From the 116 analyzed leeches, gut content information was obtained from 57 (49.13%). These 57 leeches covered the six families Glossiphoniidae, Piscicolidae, Erpobdellidae, Macrobdellidae, Haemopidae and Xerobdellidae and, within these, eight genera. Of the 57 leeches, 21 represented bloodfeeding species, 18 were liquidosomatophagous species and 18 were macrophagous species. It is important to mention that, under the classification system based on feeding preferences, seven separate leech lineages are represented. Bloodfeeding leeches are represented by three lineages: *Myzobdella, Macrobdella* and *Haementeria*+*Placobdella*. Macrophagous leeches are also represented by three separate lineages: *Erpobdella, Haemopis* and *Diestecostoma*, each lineage represents an independent transition from a bloodfeeding ancestor to macrophagous descendants (Borda & Siddall, 2004; Siddall, et al., 2016; Tessler, et al., 2018). Finally, liquidosomatophagous leeches included in this study represent a single lineage with an inferred bloodfeeding ancestor.

In addition to confirmatory sequences for the collected leeches, a total of 69 taxa were detected in the leech intestines and crops, and these spanned five phyla and 15 orders (Figure 1). These taxa are all known to inhabit the geographical area where the leeches were collected (Supplementary information) and varied in body-size (small size such as the drain fly *Psychoda alternata* and larger size such as the black-crowned night heron *Nycticorax nycticorax*), habitats (terrestrial such as earthworms of the family Lumbricidae and aquatic such as crustaceans from the family Cyclopidae), and we even detected a parasitic nematode from the family Chromadorea; note that this taxon was not considered part of the diet since it was found encysted in the gastric tissues, but is considered part of the gut content. Out of the 68 detected diet taxa (excluding the nematode), 24 could be assigned to species-level, whereas 6 could be assigned to genus-level, 6 to family-level and 4 to order-level taxa.

**Figure 1.**
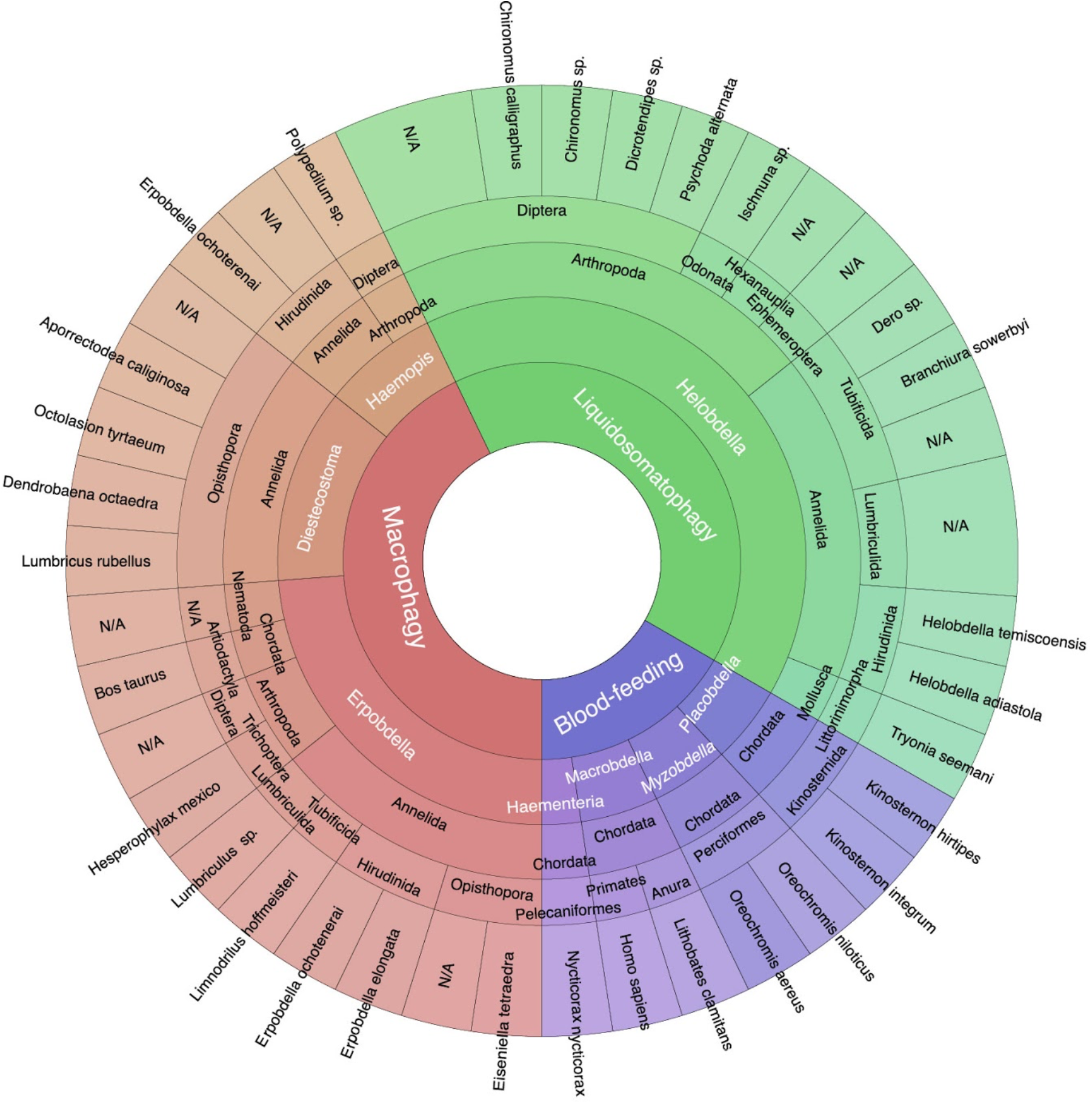
Taxa detected using DNA metabarcoding of the gut content of leeches collected in Mexico and Canada representing three feeding styles. Feeding habits of the leeches and their genus-level identifications are shown in white font, while detected dietary taxa are shown in black font. The taxonomic identification of the diet includes phylum-, order- and species-level assignment. N/A indicates that taxonomic information at order- or species-level could not be obtained. Figure created using Krona (Ondov, Bergman, & Phillippy, 2011).

The taxa detected in the 21 bloodfeeding leeches exclusively belong to the phylum Chordata without any match to invertebrate DNA. The detected chordates present different lifestyles; semi-aquatic (such as mud turtles of the genus *Kinosternon*), volant (such as the black-crowned night-heron *N. nycticorax*), amphibian (such as the bronze frog *Lithobates clamitans*) and aquatic (such as tilapias from the genus *Oreochromis*). Human DNA was detected in only one individual of *Macrobdella decora* and it probably belongs to one of the coauthors (SK), who collected this leech using his own legs. The taxa detected in the 18 liquidosomatophagous leeches were invertebrates from the phyla Annelida, Arthropoda and Mollusca, with Arthropoda and Annelida showing the highest number of taxa (eight each) (Figure 1). The identified annelids, the mollusc and the arthropods all have an aquatic lifestyle during their lifecycle. Finally, for the 18 macrophagous leeches, the taxa detected belong to the phyla Annelida, Arthropoda and Chordata; the detected annelid species could only be identified to the genus-level and has either an aquatic or terrestrial lifestyles depending on the species, the vertebrate taxon is terrestrial and the arthropods are aquatic in their larval stage. Diet identification was successful for the terrestrial leech *Diestecostoma mexicana*; note that the diet of this leech was heretofore unknown, but we can now robustly include it in the macrophagous group as only oligochaete DNA was detected. In addition to the detection of diet in the macrophagous *Erpobdella obscura*, a nematode from the family Chromadorea was detected in three individuals. In both liquidosomatophagous and macrophagous leeches, DNA from other leeches was detected, indicating at least order-level cannibalism.

### 3.2 Detection rate of diet

Diet taxa could be identified in all six analyzed leech families and within leeches spanning the three feeding modes. From the total number of analyzed leech individuals known to be bloodfeeding (33), diet taxa were detected in 21 (63.6%). Diet was detected in 53.3% (eight out of 15) of the specimens belonging to *Placobdella mexicana*, 25% (one out of four) of the specimens of *Haementeria officinalis*, 100% (10 out of 10) of the specimens of *Myzobdella patzcuarensis* and 50% (two out of four) of the specimens of *Macrobdella decora*. In the 34 analyzed liquidosomatophagous individuals, diet taxa were detected in 18 samples, corresponding to 52.9%. Of these, diet was detected in 68.7% (11 out of 16) of the specimens belonging to *Helobdella adiastola*, 50% (four out of eight) of the specimens of *Helobdella elongata*, 40% (two out of five) of the specimens belonging to *Helobdella socimulcensis* and 20% (one out of five) of the specimens of *Helobdella temiscoensis*. For the 49 macrophagous individuals, 41 known to be macrophagous and 8 *D. mexicana* with previously unknown feeding preferences, diet taxa were detected in 18 leeches (36.7%): 26.9% (seven out of 26) of the specimens belonging to *Erpobdella ochotenerai*, 40% (two out of five) of specimens of *E. obscura*, 20% (two out of 10) of specimens of *Haemopis caballeroi*, and 87.5% (seven out of eight) of specimens belonging to *D. mexicana*.

The number of vertebrate taxa detected in the 21 bloodfeeding leeches ranged from one to two. The only leech in which two vertebra taxa were detected was the bloodfeeding *M. decora* feeding on *Homo sapiens* (order Primates) and *Lithobates clamitans* (order Anura). For macrophagous and liquidosomatophagous leeches, the diet taxa ranged from one to three per individual leech; in most of the leeches only a single taxon was detected. Two invertebrate taxa were detected in the macrophagous leeches *E. ochotenerai* feeding on *Eiseniella tetraedra* (order Lumbriculida) and *Hesperophylax mexico* (order Trichoptera), in *H. caballeroi* feeding on *Polypedilum* sp. (order Diptera) and an unidentified clitellate taxon, and in *D. mexicana* feeding on *Dendrobaena octaedra* and *Lumbricus rubellus* (order Lumbriculida). The only leech species in which three invertebrate taxa were detected in the diet was the liquidosomatohagous *H. adiastola*, which was found to feed on invertebrates such as insects, oligochaets and crustaceans (Table 2).

**Table 2.**
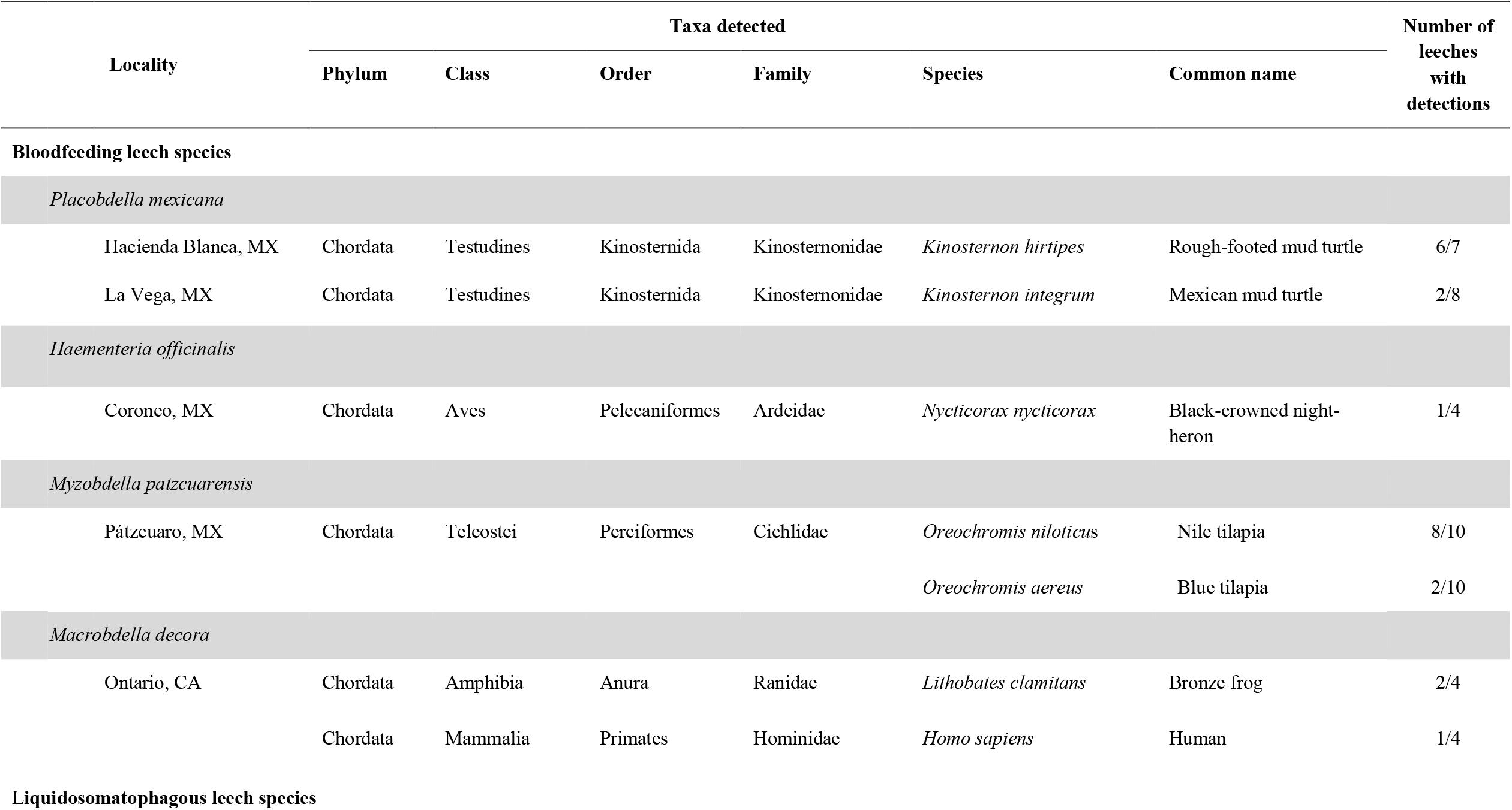
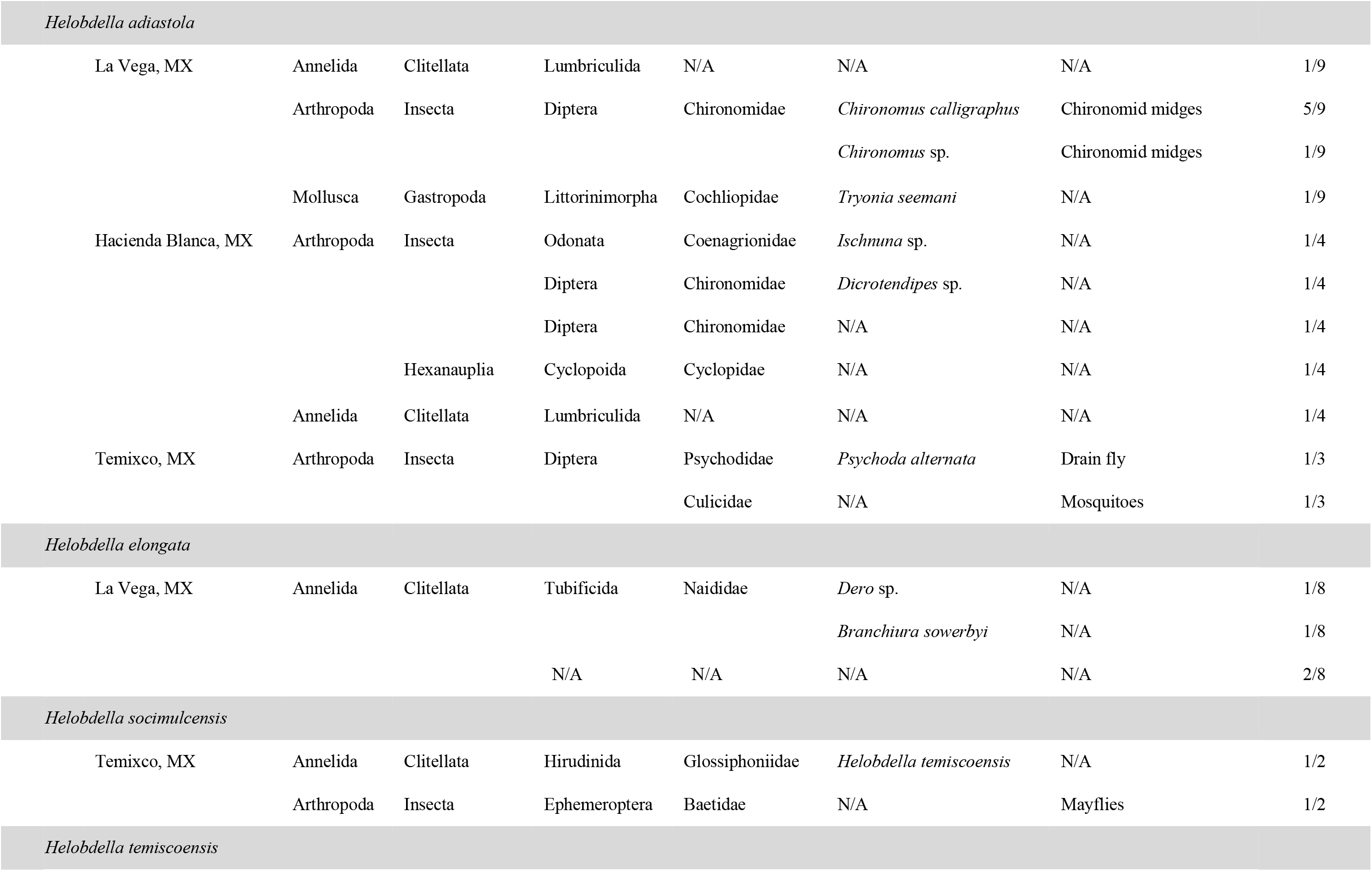
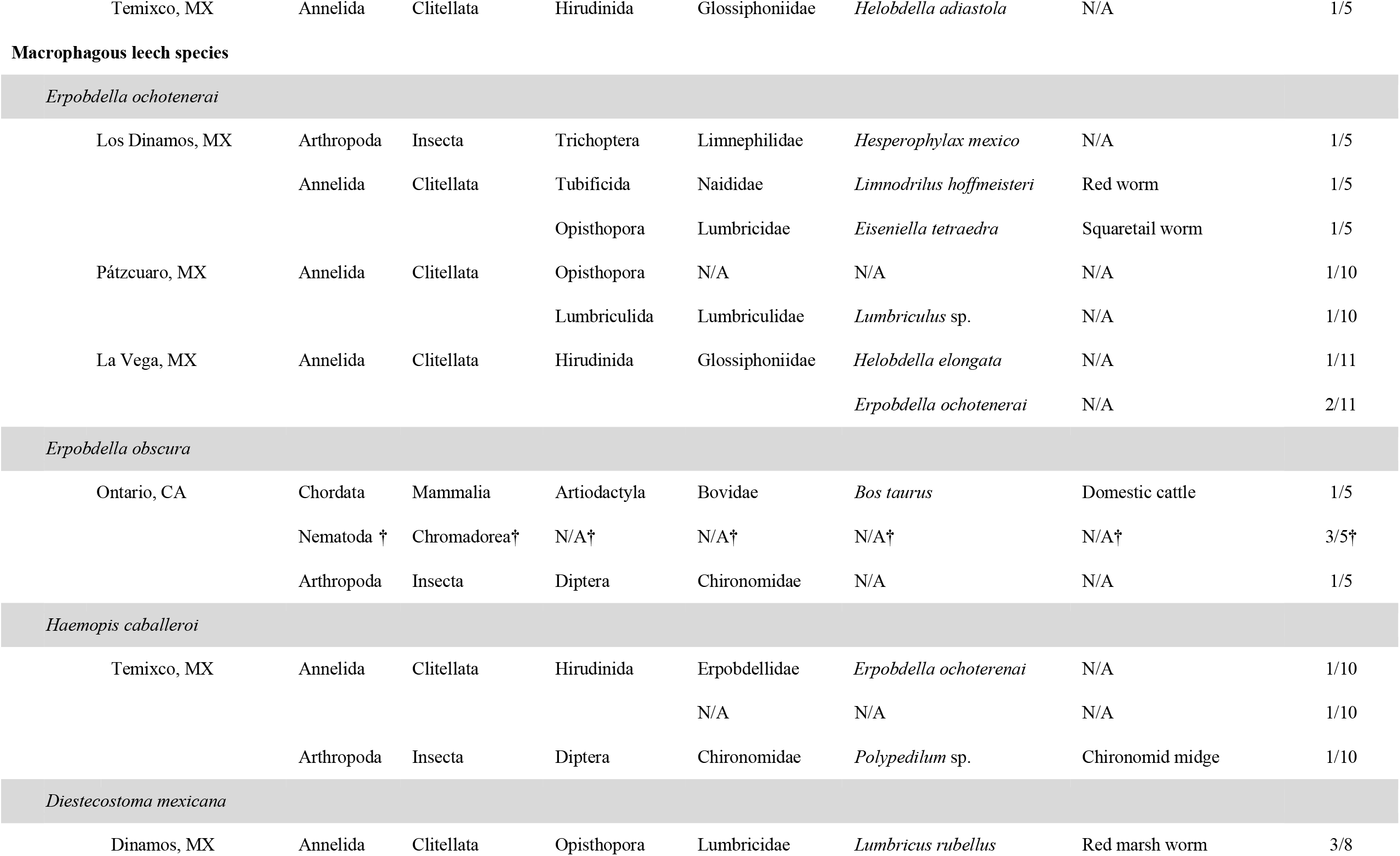
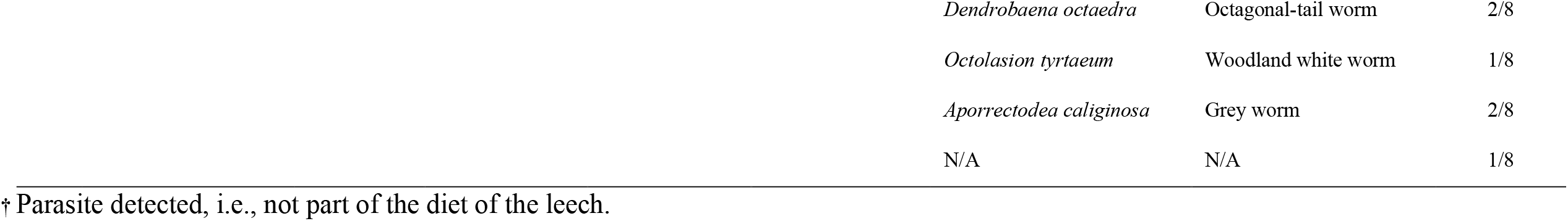
Taxa detected in the digestive tract of leeches collected in Mexico (MX) and Canada (CA), including their taxonomic classification, common name, and ratio of detection versus number of individuals analysed. Leeches are separated by known feeding habits and locality.

## 4. Discussion

In this study, we used DNA metabarcoding on iDNA from leech gut contents to obtain diet information from a broad swath of non-haemadipsid leech diversity from Canada and Mexico. The leeches encompassed eight genera from six families and the three feeding modes (hematophagy, macrophagy and liquidosomatophagy) known to be displayed by leeches. The diet taxa detected in these were aquatic and terrestrial vertebrates and invertebrates. In addition, our results show the high potential of metabarcoding using iDNA from aquatic bloodfeeding leeches to be used for vertebrate monitoring in areas with unknown animal diversity; note that the present study is the first to assess this potential in North America. In the following, we discuss the dietary range of the collected leeches within the three feeding preferences, and use that information to further discuss the utility of iDNA from leech gut contents in monitoring and estimating animal diversity.

### 4.1. Leech diet taxa

We identified the dietary range of aquatic leech groups for which detailed information is scarce, including the bloodfeeding leech species *Placobdella mexicana, Haementeria officinalis, Myzobdella patzcuarensis* and *Macrobdella decora*; the liquidosomatophagous leech species *Helobdella adiastola, Helobdella elongata, Helobdella socimulcensis* and *Helobdella temiscoensis*; and the macrophagous leech species *Erpobdella ochotenerai, Erpobdella obscura*, and *Haemopis caballeroi*. In addition, diet information was obtained from the terrestrial *Diestecostoma mexicana*, the diet of which was hitherto unknown. The low sample size for each leech species (from 4 to 26 individuals) and overall individuals per feeding habit (33 to 49) does not necessarily allow for reliable statistical analyses or for comprehensive conclusions regarding the diet of the collected leeches. Nevertheless, it does allow us to expand the knowledge about the dietary range in the different groups and to assess the potential of this tool to be used for estimating biodiversity in a region using only leech iDNA. Briefly, when comparing the diversity of diet taxa detected between the leeches with the three feeding habits, aquatic bloodfeeding leeches had the least diverse diet based solely on vertebrate blood, whereas the macrophagous leeches had the highest diversity with a diet based mostly on freshwater invertebrates (Table 2, Figure 1).

We expected to find that the diet of the four bloodfeeding aquatic leech species included only vertebrates, as previous dietary studies of both aquatic and terrestrial bloodfeeding leeches, have resulted in a vertebrate-centric diet (e.g. Drinkwater et al., 2020; Fahmy et al., 2019; Williams et al., 2020). As in Williams et al., (2020), we detected both aquatic and terrestrial vertebrate taxa in the analyzed aquatic bloodfeeding leeches (Table 2; Figure 1). Unsurprisingly, this is in contrast to the well-studied, terrestrial, bloodfeeding haemadipsid leeches, in which no aquatic vertebrates have so far been detected (Drinkwater et al., 2020, 2018; Hanya et al., 2019; Schnell et al., 2018; Schnell et al., 2012). Among the terrestrial vertebrates that we detected in the aquatic bloodfeeding leech species, we identified a single bird species, the black-crowned night heron, *N. nycticorax*. This bird species was detected in a leech belonging to the genus *Haementeria*, which is known to generally feed on mammals and reptiles (Charruau et al., 2020; Oceguera-Figueroa, 2012). As this bird was detected in only one individual leech, more sampling is needed to determine whether *Haementeria officinalis* commonly feeds on birds. Birds have been involved in leech passive and active dispersal (Davies et al., 1982; Siddall et al., 2013) and our findings suggest that the black-crowned night heron may be involved in the dispersion of this leech along its broad geographical distribution in central Mexico. Other vertebrates detected in the diet of aquatic bloodfeeding leeches included invasive aquatic species. In the diet of the aquatic bloodfeeding leech *Myzobdella patzcuarensis* we detected the Nile tilapia and the Blue tilapia, both of which were introduced to the lake of Pátzcuaro for the purpose of breeding consumption fishes (Berlanga et al., 1997). *Myzobdella patzcuarensis* is known to be a parasite of fish, and has previously been found to feed on both native and exotic fish species (López-Jiménez, 1985), and this generalist pattern is further corroborated by our findings. Finally, both mammal and amphibian DNA were detected in the diet of the bloodfeeding *Macrobdella decora*. Despite the fact that metabarcoding cannot distinguish between life stages of prey (i.e., eggs or adults), our finding of frog DNA inside this leech, may reinforce previous reports that it occasionally ingests frog eggs (Moore, 1953; Trauth & Neal, 2004; Turbeville & Briggler, 2003). Leeches with a liquidosomatophagous diet were found to feed exclusively on invertebrates (Figure 1). We found that members of the liquidosomatophagous genus *Helobdella* suck molluscs, arthropods (insects and crustaceans) and clitellates (oligochaetes and hirudineans) (Figure 1). However, despite that we observed molluscs in the three locations where the *Helobdella* individuals were collected, only one mollusc taxon was detected in a single leech. The prevalence of insects in the leech diet was therefore surprisingly high based on previous observations, as it is known that the liquidosomatophagous leeches feed mainly on molluscs and oligochaetes (Sawyer, 1986). Notably, two individuals of *Helobdella* were found to have fed on another congeneric hirudinean. This has been previously recorded as a rare behaviour in *Helobdella fusca* (Mathers, 1948), but our results indicate that cannibalism may be more widespread than previously thought for the genus *Helobdella*, with *H. temiscoensis* and *H. socimulcensis* also presenting this behaviour. However, as this was only found in two individuals, sampling of more species is needed to fully understand this behaviour.

Macrophagous leeches were found feeding on invertebrates, which corresponds to previous knowledge about the diet of leeches with this feeding habit (Sawyer, 1986). There was a clear preference for an annelid-based diet in the analyzed leeches, as 13 out of the 17 taxa detected in the gut contents were annelids, including some hirudineans. The direct observation of the same species of leech inside the digestive tract during dissection supports this metabarcoding results. In addition, cattle DNA was also detected in one individual of the genus *Erpobdella*. Although Kutchera (2003) previously recorded this leech sucking body fluids from dead decaying vertebrates, the detected cattle could potentially be a contamination by the bovine serum albumin (BSA) used for the PCR amplification. Therefore, analysis of more individuals is needed to clarify if this is a common behaviour in the species of the genus *Erpobdella*.

The diet of the terrestrial leech *Diestecostoma mexicana* was previously unknown, but it is expected to be a bloodfeeder as it coexists with salamanders (Caballero, 1940). Our results indicate that this is not the case, as the eight individuals of *D. mexicana* were found to feed exclusively on oligochaetes, with a total of five taxa detected in their gut contents (Table 2, Figure 1). This is the first study to attempt to solve this issue, and this leech can now be inferred to be macrophagous.

### 4.2 Non-haemadipsid leeches for animal monitoring

The detection of DNA from turtles, fish and birds, provides different opportunities to use non-haemadipsid leeches as a monitoring tool. Given that iDNA from leech gut contents can be used as a tool to monitor e.g. otherwise elusive and endangered vertebrates (Nguyen et al., 2021; Schnell et al., 2012), it is surprising that non-haemadipsid leeches have been almost completely neglected for this purpose (Williams et al., 2020).

As in previous iDNA studies of leech gut contents (Schnell et al., 2018), we detected birds, reptiles, and amphibians in the aquatic bloodfeeding leeches; note that reptile DNA has not been previously detected in this group of leeches (Williams et al., 2020). However, with the exception of human DNA, we did not detect other mammals, despite this being a commonly targeted group for iDNA studies. Nevertheless, and importantly, we did detect two fish species, both of which are known to be invasive species. Animal monitoring is commonly used not only to detect the presence of rare species but also the presence of invasive species, as they may represent a serious challenge to the health of ecosystems (Ota, Hall, Malloy, & Clark, 2020). In aquatic ecosystems, invasive fish can have extreme negative consequences on biodiversity and therefore their monitoring is of vital importance (Trebitz et al., 2017). Using molecular methods to monitor fish minimizes the onus on taxonomic expertise, in contrast to using traditional methods such as fishing (Bessey et al., 2020). Through such an approach, aquatic bloodfeeding leeches from the family Piscicolidae, e.g. *Myzobdella patzcuarensis*, can be leveraged to monitor fish faunal changes in aquatic environments, for both native and invasive fish species. Importantly, other types of samples, such as environmental DNA (eDNA) from water, can provide a complete overview of the aquatic communities (Cantera et al., 2019) and can therefore be a more viable option.

Whereas iDNA studies have traditionally targeted vertebrate DNA, our study shows that invertebrate DNA is equally viable. This is the case for macrophagous and liquidosomatophagous leeches. Knowledge of the presence of certain arthropods or annelids, which are the taxa most commonly found in the guts of these leeches, can provide important information. For example, invasive earthworms are known to be a menace to several freshwater ecosystems (Ziemba et al., 2016). Annelid-eating macrophagous leeches such as those in the genera *Haemopis, Diestecostoma* and *Erpobdella*, can be used to monitor earthworms, especially in areas where invasive species are found. It is important to mention that even though a group of leeches is known to feed on a certain animal group, the diet is often more flexible. For example, although the liquidosomatophagous genus *Helobdella* was only found to feed on invertebrates, other studies have found the leech to be a facultative parasite of amphibians (Tiberti & Gentilli, 2010; Zimić, 2015) and fish (Malek & McCallister, 1984). Therefore, if *Helobdella* is collected for iDNA studies, there is the potential for retrieving not only invertebrate but also vertebrate information. Our results provide proof-of-concept that leeches exhibiting all three feeding modes can provide information about which animal taxa live in the same area, and that iDNA from leech gut contents can be used in other geographical areas outside the Indo-Pacific region.

The ease with which samples can be collected is important to take into account when using bloodfeeding organisms as biomonitoring tools. Terrestrial bloodfeeding leeches are relatively easy to collect as they readily parasitise humans and will therefore actively hunt the collector (Schnell et al., 2012). Similarly, the collection of aquatic leeches is relatively straightforward, even when targeting non-bloodfeeding leeches. Several leeches can be found under rocks, plants and debris submerged in water. In addition, if submerging the collector’s legs at the same time, bloodfeeding leeches can also be collected from the exposed skin. Contrary to this, the terrestrial macrophagous leech, *Diestecostoma mexicana*, was not easily collected, and many hours were spent trying to find only a few individuals. Therefore, if the aim is to use a group of leeches for regular animal monitoring, an easily collected group should be chosen, such as the aquatic leeches; specifically bloodfeeding species if targeting vertebrates that live in or visit in a specific waterbody, or liquidosomatophagous and macrophagous leeches if targeting small invertebrates. It is also important to ensure that the collection of leeches for iDNA studies will not endanger the leech populations. In this study, none of the collected leeches are threatened and can therefore continue to be collected for diet analyses. Note here that the two commonly used European medicinal leech species, *Hirudo medicinalis* Linnaeus, 1758 and *Hirudo verbana* Carena, 1820 are both on the IUCN red list and should not be collected (see Williams et al., 2020).

Further studies focusing on animal diversity, especially in aquatic environments, should compare the use of other methods, e.g. water or soil eDNA, camera traps, fishing, and iDNA from leech gut contents, to clarify if these methods are comparable or complementary. We hypothesize that, as seen in other iDNA studies (Abrams et al., 2019; Tilker et al., 2020; Weiskopf et al., 2018), the use of these leeches for animal monitoring may not provide a complete overview of the local fauna but may provide complementary information to other monitoring methods.

### 4.3 Technical considerations

In this study, a universal primer set targeting a partial fragment of the COI barcode region (mlCOIintF/jgHCO2198) (Leray et al., 2013; Geller et al., 2013) was chosen to allow detection of diet across a broad taxa range of metazoans (Leray et al., 2013). In addition, this primer set can also provide information about possible metazoan endoparasites (Bohmann et al., 2018) and can simultaneously co-amplify leech DNA, thereby providing DNA barcode-based host identification. Nevertheless, this co-amplification can cause lower detection rates of the diet, as the leech DNA can overpower the diet DNA in the leech gut contents. To ensure that the diet is also detected, the screening of samples using qPCR prior to tagged PCR is important, as it increases the detection probability of DNA in both high and low copy numbers (Murray et al., 2015).

Based on our results, future studies can easily gauge the targeted iDNA by employing more or less specific primer sets to amplify for a chosen taxon. However, it is important to keep in mind that some leeches also feed on other hirudineans. In the present study, this was found for the genera *Erpobdella, Helobdella* and *Haemopis*, and has also been recorded in other leeches previously (Aminov, 2019). The ingestion of leeches from a different genus can be easily detected with molecular methods using primer sets that have been shown to scrutinize at species-levels, but the ingestion of a member of the same species may still pose issues. Unless direct observations of this have been made, e.g. in *Erpobdella* as in the present study, it is difficult to assess if the DNA detected is from the leech or from the diet. As has been shown for arthropods (Elbrecht et al., 2018) and recently with rabbit DNA within iDNA of leeches (Nguyen et al., 2021), one way to overcome this is to determine if the leech and diet haplotypes are fully conserved; if a certain haplotype can be evinced from non-gut tissue then seperate haplotypes within the ingesta can infer species-level cannibalism.

Processing the leeches individually allowed us to have a better overview of the detection rate of animal DNA. With the exception of Hanya et al., (2019) who extracted and PCR amplified the DNA of leeches individually, most studies using DNA metabarcoding to target iDNA from leech gut contents have processed the leeches in pools of several individuals (Abrams et al., 2019; Fahmy et al., 2019; Williams et al., 2020). In addition, except for Hanya et al., (2019), Schnell et al., (2012) and Schnell et al., (2018), all previous studies, have used several metabarcoding primer sets and, therefore, a direct comparison to our results is strenuous, as we processed the leeches individually and used only a single metabarcoding primer set. In addition, no other study has used metabarcoding to target iDNA from the gut contents of non-bloodfeeding leeches. Nevertheless, in some ways, it is possible to compare our results to those of previous studies. Studies pooling several leeches have found between 0.5 to 1.6 diet taxa per pool (Abrams et al., 2019; Axtner et al., 2019; Drinkwater et al., 2020; Schnell et al., 2018; Williams et al., 2020), whereas Hanya et al., (2019) and Schnell et al., (2012) detected 1-3 diet taxa (including human) in 24.3% of their individual leeches. In the current study, diet taxa were detected in more than 50% of the individually analyzed bloodfeeding and liquidosomatophagous leeches, whereas the detection rate in the macrophagous leeches was 36.7%. Despite this lower detection, two diet taxa were detected in three macrophagous leeches, while in the other two groups the majority of detections were of one taxon per leech. Our higher detection rate can be due to the way leeches were collected, as many of them we found under objects submerged in water which may indicate that they were not seeking food; however, this could be also be the result of the bloodfeeding *M. patzcuarensis* being collected directly from its host, thus increasing the probability of detecting fish DNA. Moreover, the processing of the samples can also contribute to the paucity of diversity in any individual leech. Hanya et al., (2019) reported variations in the detection of mammal species when the DNA extract also contained leech DNA and when using different primer sets. This could be another reason for the higher detection rate in our study as we did not pool leech specimens. Likewise, the degradation rate of the ingested DNA can also play an important part in the diet detection. Currently, it is still unknown if DNA from the diet of liquidosomatophagous and macrophagous leeches is preserved for at least four months within the intestine, as it is in bloodfeeding leeches (Schnell et al., 2012). Future studies should address this.

Finally, it is important to remember that the strategy employed by the present study is based on DNA barcodes and, thus, the lack of a robust and well-curated comparative database will negatively impact the potential for taxonomic identification of the obtained sequences. In the present study, 11 out of the 40 OTUs detected could not be identified to species level, therefore providing much less informative results. As databases become more complete, so will the taxonomic assignment of the detected taxa and, with this, our knowledge of the diet of this group of invertebrates will dramatically increase.

## 5. Acknowledgments

Leeches were collected under the scientific collecting permits SEMARNAT (Mexico), issued to AOF. CL was supported by the National Council of Science and Technology in Mexico (CONACyT) Grant CVU 582667 and the Danish Council for Independent Research, DFF grant 7079-00029B.

## 6. Author contribution

CL, AOF, KB and MTPG designed the study. CL, AOF and SK collected the samples. CL generated and analysed the data. CL wrote the original manuscript, with input and revisions from all the co-authors.

## 7. Data accessibility

All sequenced data and tag information will be available on the University of Copenhagen Electronic Research Data Archive (UCPH ERDA) Digital Repository, DOI: xxx upon acceptance of the manuscript.

