## Supplementary Information for "The potential of aquatic haematophagous, liquidosomatophagous and macrophagous leeches as a tool for iDNA characterisation"

**Supplementaray Methods**

Leeches collected in Mexico are from the following locations: Hacienda Blanca in the state of Querétaro (2280 masl; 20°15'44.3"N 100°04'25.3"W), La Vega in the state of Jalisco (1260 masl; 20°35'39"N 103°50'48"W), Coroneo in the state of Guanajuato (2380 masl; 20°17'58"N 100°25'44" W), Temixco in the state of Morelos (2240 masl; 18°51'30"N 99°13'32" W), Pátzcuaro in the state of Michoacán (230 masl; 19°32'29"N 101°38'26"W) and Los Dinamos in Mexico City (2960 masl; 19°18'5"N 99°18'54" W).
Leeches collected in Canada are from the following locations: *Erpobdella obscura,* unnamed pond from west side of Brent Rd, NW of Windigo Lake (46°09'19.7"N 78°19'15.0"W); *Macrobdella decora,* Clear Lake, Grundy Provincial Park (45°55'56.9"N 80°34'29.9"W).

The geographic distribution information of the taxa was retrieved from different sources: The Reptile Database (https://reptile-database.reptarium.cz/), the Global Biodiversity Information Facility (https://www.gbif.org), iNaturalist (https://www.inaturalist.org), Enciclo vida (https://enciclovida.mx) and Czaja et al (2020).

**Supplementary Results**

**Supplementary Table1.** Number of reads after each filtering step.

| | Raw sequences | After adapter remov and merging | After begum sort |
|---|---|---|---|
| **Pool1** | 1,676,418 | 932,053 | 811,597 |
| **Pool2** | 4,467,638 | 3,825,847 | 2,744,200 |
| **Pool3** | 4,065,352 | 3,754,973 | 2,825,769 |
